# Periodic spatial patterning with a single morphogen

**DOI:** 10.1101/2022.03.21.484932

**Authors:** Sheng Wang, Jordi Garcia-Ojalvo, Michael B. Elowitz

## Abstract

Multicellular development employs periodic spatial patterning to generate repetitive structures such as digits, vertebrae, and teeth. Turing patterning has long provided a key paradigm for understanding such systems. The simplest Turing systems are believed to require at least two signals, or morphogens, that diffuse and react to spontaneously generate periodic patterns. Here, using mathematical modeling, we show that a minimal circuit comprising an intracellular positive feedback loop and a single diffusible morphogen is sufficient to generate stable, long-range spatially periodic cellular patterns. The model considers cells as discrete entities as a key feature, and incorporates transient boundary conditions. Linear stability analysis reveals that this single-morphogen Turing circuit can support a broad range of spatial wavelengths, including fine-grain patterns similar to those generated by classic lateral inhibition systems. Further, signals emanating from a boundary can initiate and stabilize propagating modes with a well-defined spatial wavelength. Once formed, patterns are self-sustaining and robust to noise. Finally, while noise can disrupt patterning in pre-patterned regions, its disruptive effect can be overcome by a bistable intracellular circuit loop, or by considering patterning in the context of growing tissue. Together, these results show that a single morphogen can be sufficient for robust spatial pattern formation, and should provide a foundation for engineering pattern formation in the emerging field of synthetic developmental biology.

## Introduction

The fundamental question of how periodic spatial patterns emerge in multicellular organisms arises both in the context of natural developmental systems and in the emerging field of synthetic development, where researchers seek to engineer biological circuits that recapitulate fundamental aspects of multicellular development (Davies 2008; Ebrahimkhani and Ebisuya 2019; Toda et al. 2018). Understanding the minimal requirements to generate spatial patterning could provide insight into natural patterning processes, and offer a basis for engineering synthetic patterning circuits with minimal complexity.

Previous work has focused on three general paradigms for spatial periodicity (Schweisguth and Corson 2019). First, studies of early embryonic development in *Drosophila* unveiled a far-reaching scenario for patterning based on positional information and hierarchical morphogen gradients (Wieschaus 2016; Briscoe and Small 2015). Second, at the other extreme, classic Turing patterning systems use interactions between two morphogens to spontaneously generate periodic spatial patterns (Kondo and Miura 2010). Finally, short-range alternating patterns can be produced by lateral inhibition mechanisms in which neighboring cells establish opposite fates through direct signalings, like those implemented by the Notch pathway (Sprinzak et al. 2011; Corson et al. 2017; Collier et al. 1996). While powerful, these three mechanisms may not, individually, be ideal for all purposes or may be more complex than necessary. For example, positional information-based mechanisms do not readily scale to pattern an arbitrarily extended region (Ben-Zvi, Shilo, and Barkai 2011), potentially limiting their evolvability. In turn, classic Turing mechanisms require multiple morphogens and fine-tuning of key parameters (Marcon et al. 2016; Scholes et al. 2019), adding complexity for synthetic applications. Finally, lateral inhibition patterns generally operate at small spatial length scales, comparable to that of the individual cell (Cohen et al. 2010; Hadjivasiliou, Hunter, and Baum 2016). Natural systems undoubtedly combine these and other mechanisms to achieve more complex pattern formation capabilities (Green and Sharpe 2015).

In natural contexts, periodic patterns are generated by a combination of (1) reaction-diffusion circuits operating within the developing tissue and (2) triggering signals that emerge from tissue boundaries. For example, digit patterning in the developing limb bud is set off by signals coming from the apical epidermal ridge (Raspopovic et al. 2014). Similarly, tooth patterning is thought to involve sequential activation of new primary enamel knots, through a mechanism that may involve both local morphogen dynamics and inputs from adjacent tissue (Kavanagh, Evans, and Jernvall 2007). Feather patterning appears to similarly combine these mechanisms (Kavanagh, Evans, and Jernvall 2007; Ho et al. 2019). These examples provoke the question of what minimal circuits are sufficient for periodic boundary-triggered patterning.

An ideal pattern-forming system would have several key characteristics: First, it should be triggered by signals from a localized adjacent region or boundary, to enable control of patterning. Second, the pattern would spatially propagate to fill an extended region of tissue. Third, to enable synthetic engineering, the system should be as simple as possible, utilizing a minimal number of components and interactions. The ideal system should further be robust to unavoidable fluctuations in cellular components.

Here, we introduce a remarkably simple mechanism for boundary-triggered spatial periodic patterns that uses a single morphogen. This architecture differs from classical Turing systems in that the activator is confined to the cell in which it is synthesized, and cannot diffuse to other cells. It thus represents the limit of a classic Turing system when the mobility of the activator approaches 0. Unlike Turing systems, this circuit cannot, by itself, spontaneously produce consistent, well-defined long-wavelength patterns. However, when stimulated by an adjacent signal from a localized region, or boundary, it can produce spatially propagating periodic patterns in the levels of its components that self-sustain even after the initiating signal is removed. This single morphogen mechanism demonstrates how complex spatio-temporal patterns can arise from remarkably simple underlying circuit mechanisms, and provides a realistic design for synthetic developmental patterning systems.

## Results

### A single morphogen is sufficient to form patterns on the spatial discrete lattice

Alan Turing’s patterning circuit in his 1952 paper is one of the mathematically simplest models that can form patterns through diffusion-driven instability (Turing 1952). The Turing circuit comprises two components that regulate their own production and/or degradation, and diffuse as morphogens (Figure 1A, upper left panel). Patterning occurs when the inhibitory morphogen has a larger diffusion coefficient than the activating morphogen, causing local activation and long-range inhibition (Figure 1A, left row) (Murray 2001).

**Figure 1:**
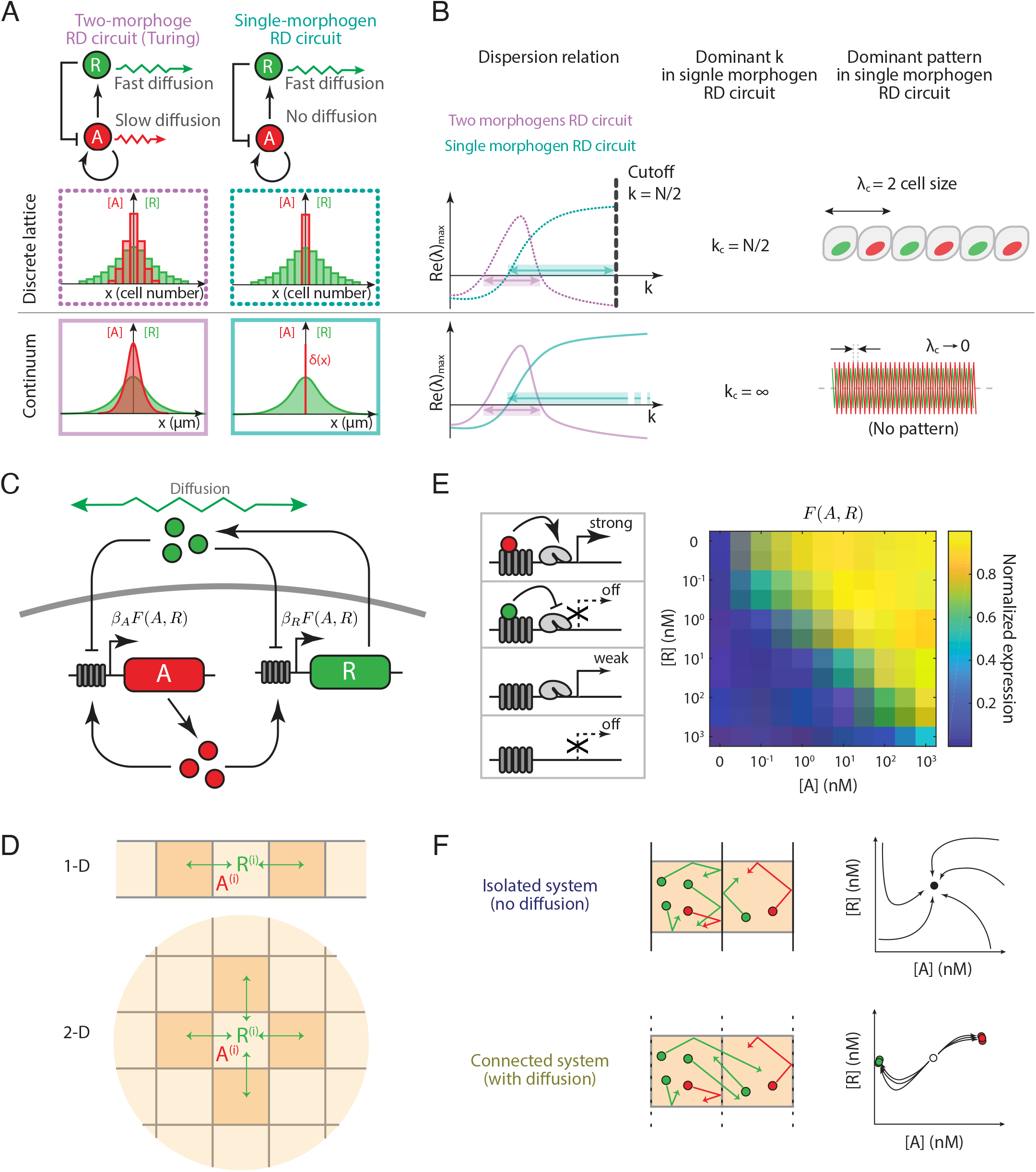
The lattice ODE simulation of a minimal single-morphogen reaction-diffusion circuit evaluates the pattern formation mechanism in discrete space. (A) **The single-morphogen reaction-diffusion circuit obeys the “local-activation and long-range inhibition” rule on the discrete lattice.** Classical Turing circuit requires at least two diffusive species, while in the single-morphogen reaction-diffusion circuit, the repressor is the only component that could diffuse among the cells. In specific parameter regimes, the classical Turing mechanism could form patterns on both continuum and discrete lattice, though the power frequency selection is different in these two cases (Plahte 2001). The patterning on discrete lattice provides the activator with at least two-cell size spatial period. (B) **The shortest wavelength on the discrete lattice determines the upper bound of the wavenumber for the single-morphogen reaction-diffusion circuit.** Stability analysis reveals that once the SMRD circuit drives the patterning, the dominated wavenumber of the final pattern goes to infinity. This type of pattern, named “Type II Turing pattern” (Scholes et al. 2019), is purely a mathematical concept with no actual pattern in the physical world. Meanwhile, patterns on the discrete lattice intrinsically have the shortest wavelength as two-cell size, which sets a cutoff for wavenumbers and enables the reaction-diffusion patterning with a single morphogen. (C) **The minimal SMRD circuit contains two components driven by the same promoter, a non-diffusive activator upregulating both production and a diffusive repressor downregulating the production.** (D) **The simulation executes on a chain (1-D) or a square lattice (2-D), where each lattice node directly contacts two or four neighboring nodes, respectively.** The isolated system forbids diffusion of R among cells so that each lattice node is well separated. Meanwhile, the connect system allows free diffusion of R. i denotes the cell counts number in the lattice. (E) **The promoter integrates the upregulation from A and the downregulation from R in a ratiometric fashion.** The model assumes four possible promoter binding states, polymerase + activator, polymerase + repressor, polymerase only, and empty cassette. The portion of the four states shaped by the A and R concentrations determines the promoter activity F(A, R). The calculated titration matrix reveals that the promoter activity has a ratiometric relation of co-existing A and R. (F) **The simulation starts from the homogeneous state and ends up with stable patterns.** In the isolated system where no diffusion is allowed among cells, the lattice stays at its homogeneous steady-state. In the connected system, if a circuit is patternable, this steady-state should become unstable so that cells would bifurcate and evolve into heterogeneous patterns. Meanwhile, a non-patternable circuit is still stable under the connected system.

Since there is no lower bound for the diffusion coefficient of the activating morphogen, we wondered whether the system could support pattern formation in the limit of no activator diffusion at all, i.e. if the activator were confined to a single cell rather than diffusing as a morphogen. In continuous systems, the limit of no activator diffusion does not pattern due to the unbounded spectrum of high-frequency modes (Figure 1A, lower right panel). However, on a discrete lattice, cells expressing the non-diffusive activator exhibit local (same-cell) activation, while inhibiting cells in their extended neighborhood (Figure 1A, upper right panel). In this case, self-activation occurs on the scale of a single cell, providing a high-frequency cutoff.

To understand whether such a discrete system could form robust patterns and, if so, what characteristics these patterns might have, we performed linear stability analysis of both discrete and continuous single-morphogen systems (Murray 2001). We analytically computed the spatially homogeneous steady state of the system, and analyzed its stability to harmonic perturbations with varying wavenumber k, by determining the eigenvalue of the Jacobian of the system with largest real part, and its dependence on k (dispersion relation) (Murray 2001). In a classic two-morphogen Turing system, the real part of that eigenvalue could take on a maximum value at an intermediate wavenumber (Figure 1B, left panels, purple lines). By contrast, in a single-morphogen system, the resulting dispersion relation increases monotonically with increasing k (decreasing spatial period), as shown by the blue lines in the left panels of Figure 1B. This monotonic increase suggests that the resulting pattern should be dominated by perturbations with arbitrarily large spatial frequencies. In the continuous limit, this situation leads to physiologically unreasonable patterns, which involve infinitely small wavelengths (Figure 1B, lower right panel). However, a discrete lattice imposes a cutoff at a wavelength of two cell diameters (Figure 1B, upper left panel), and can thereby support patterns involving alternating cell states (Figure 1B, upper right panel). Thus, a single-morphogen circuit that would not pattern in the continuous limit, could produce “fine-grain” alternating patterns on a lattice, similar to lateral inhibition patterns(Collier et al. 1996). Further, as we will see below, certain conditions, including particular initial perturbations, can further expand the spectrum of patterning behaviors to include longer wavelength patterns.

To gain more insight into particular realizations of single-morphogen patterning systems, we considered a minimal, two-component regulatory system that could be constructed synthetically in a living cell using known genes and regulatory sequences (Figure 1C). In this system, an intracellular activator denoted A, promotes its own production and that of a repressor, denoted R. The repressor, but not the activator, is secreted and can diffuse from one cell to another to down-regulate production of both A and R in both its own cell and neighboring cells. To simulate dynamics on a discrete cell lattice, we used ordinary differential equations (ODEs) to represent the dynamics within each cell, allowing R molecules to transfer from one cell to its neighbors (Figure 1D). With these assumptions, we obtain the following equations for the dynamics of the single-morphogen system on a discrete lattice:

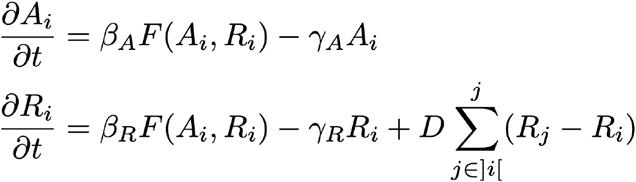

Here, β_A_ and β_R_ denote the maximal production rate of the activator and the repressor, respectively. γ_A_ and γ_R_ denote effective protein degradation and loss rates. D denotes the diffusion coefficient of the repressor. ]i[ denotes the set of cells that directly contact cell i. Finally, to mathematically represent combinatorial regulation by the activator and repressor, we assume that the two regulators compete to bind to the same target DNA binding site, a relatively simple type of interaction that can be synthetically engineered (Phillips et al. 2012; Zhu et al. 2021). This results in the following dependence of target promoter activity on A and R concentration (Figure 1E):

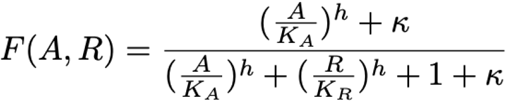

Here, κ quantifies the basal expression in the absence of activator and repressor, K_A_ and K_R_ respectively represent the concentrations required for half-maximal activation and repression, and *h* is a Hill coefficient that quantifies the ultrasensitivity of the response to either regulator. This production function F(A, R) is assumed to affect the expression of both A and R. We note that the alternative Gierer–Meinhardt reaction-diffusion kinetics, which is widely applied to model biological patterning formation, lead to similar results (see Supplementary Figure S1) (Meinhardt and Gierer 1974).

A typical reaction-diffusion simulation starts from a homogeneous initial condition in which all cells are at the same value, equal to the steady-state for the case of no diffusion, D = 0 (Figure 1F, left), perturbed by small spatial fluctuations. As time progresses, diffusion can, under some circumstances, destabilize the spatially homogeneous initial state, leading to patterning. The simulation continues until the concentration profiles of all cells reach new steady-state values (Figure 1F, right).

We first simulated a square two-dimensional lattice, starting with a low level of random noise in the initial state (Figure 2A, Movie S1). Concentrations of activator and repressor began diverging in different cells, eventually reaching two distinct states with high and low concentrations of A (Figure 2A). Spatially, these states formed a disordered “salt and pepper” pattern, with a wavelength of about 2 cell diameters, as expected from the dispersion relation (Figures 1B, 2B). Autocorrelation analysis showed that the diffusion coefficient defines the length scale of the final pattern (Figure 2C). In the low diffusion regime, the circuit generates checkerboard-like patterns (Figure 2C, upper two panels), while high diffusion rates produce irregular, low wavenumber patches of elevated A expression (Figure 2C, lower two panels). Further frequency power spectrum analysis confirmed the two regimes corresponding to the checkerboard-like pattern and less regular patch pattern (Figure 2D). Together, these results show that the discrete single-morphogen system can spontaneously pattern with a diffusion-dependent wavelength.

**Figure 2:**
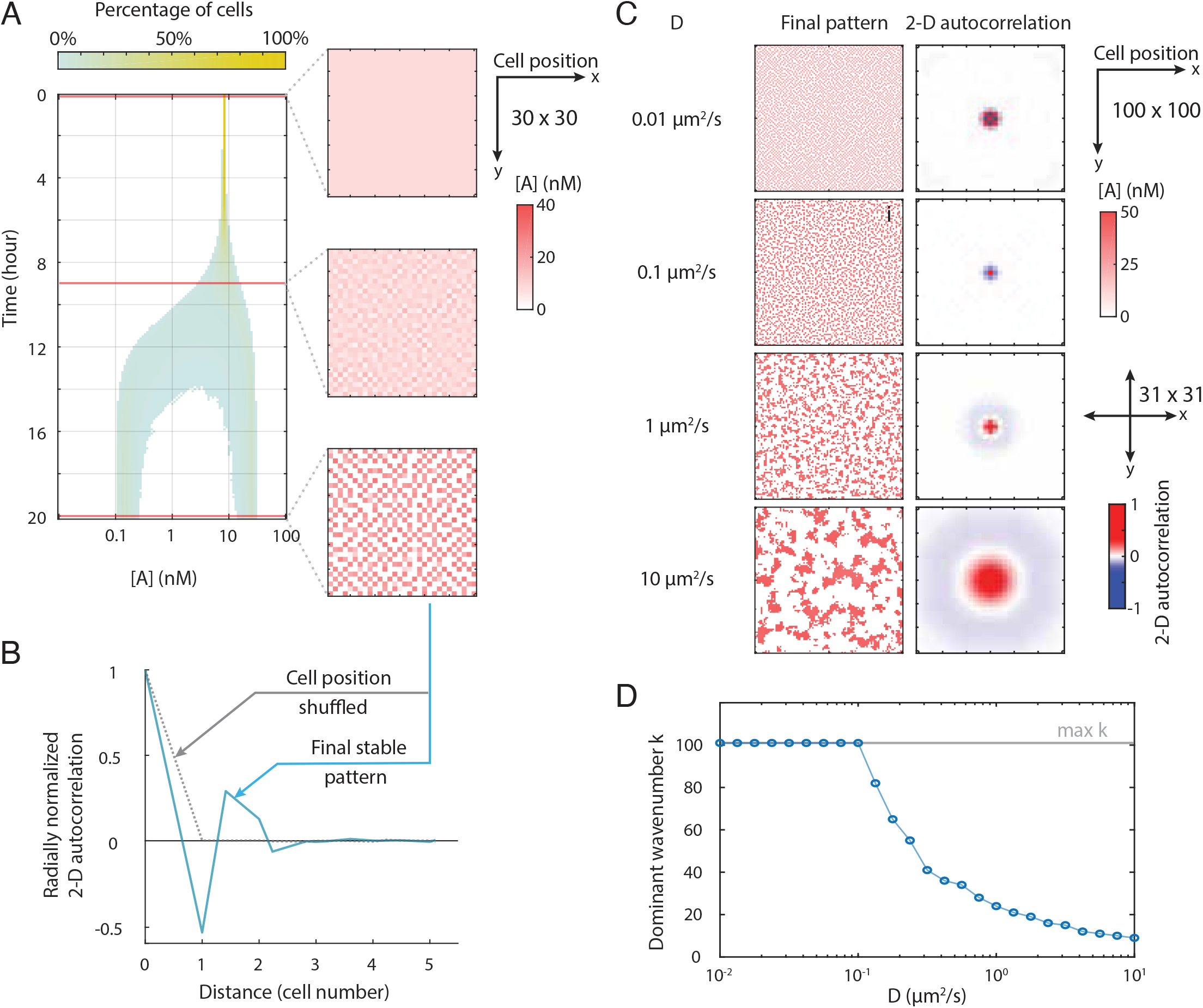
The SMRD circuit develops lateral inhibition patterns with global random noise as initial conditions. (A) **The identical lattice cells bifurcate during the simulation forming two distinct classes.** The histogram of cell count percentage based on the activator concentration captures the dynamics of the bifurcation. The lattice system finally converges to a “salt and pepper” pattern. (B) **The final heterogeneous pattern is spatially autocorrelated**. The blue curve presents the radially averaged 2-D autocorrelation of the final frame. The negative value at one cell distance reveals the lateral inhibition effect. As the control, random shuffling cell positions of the final pattern lose the spatial correlation (red curve). (C) **The diffusion coefficient of the repressor alters the range of the lateral inhibition zone.** If the repressor diffuses slowly, the lateral inhibition intensively represses the adjacent cells. Meanwhile, with a large D, the adjacent neighbors show a positive correlation, and the further neighboring cells are repressed. (D) **The dominant wavenumber in final patterns shows two types of dependence in two regimes of the diffusion coefficient.** Systems with lower diffusion coefficients produce checkerboard patterns with a constant dominant wavenumber. In the high diffusion regime, however, the final dominant wavenumber goes low when the diffusion coefficient increases.

### The single-morphogen patterning circuit can generate stable, long-wavelength periodic patterns

Given the large range of wavenumbers for which the dispersion relation is positive, one could expect patterns with low wavenumbers to be stable, as long as they fall within the above-mentioned range of positive dispersion relation. We thus asked whether initial conditions could stabilize particular low wavenumber modes, and thereby establish well-defined spatial patterns. To that end, we first investigated the effect of point and line morphogen sources (Figure 3A, left). We simulated the same system (Figure 2A) as above, but transiently increased the concentration of R to a higher value either in a single cell or a line of cells. These perturbations generated, respectively, either concentric rings or periodic lines of elevated activator expression (Figure 3A, right). The patterns exhibited several key features: First, they formed sequentially, as a front dynamically propagates away from the perturbation at a constant velocity, as shown in Figure 3B (orange arrow) and Movie S2 for the case of the line source (limitations on pattern propagation are discussed below). Second, in the final pattern, A and R exhibit in-phase periodic spatial oscillations (Figure 3C), as expected given that they are co-regulated by the same inputs (Figure 1C). Third, in contrast to standard Turing patterns in which spots of high activator concentration tend to rearrange to maximize distances from neighboring peaks, in this case, peak morphogen positions remained fixed throughout patterning (Figure 3B). Finally, once triggered, the dynamic patterning process is self-sustaining and continues to propagate even if the initiating perturbation is transient (Figure 3B, purple arrow). All of these features can occur with a variety of perturbations, including reducing R, or increasing or decreasing A (Figure S2). Even perturbations whose amplitude is small compared to the initial state concentrations can be sufficient to generate the patterns in a deterministic setting (Figure S2). Together, these results indicate that transient spatially structured perturbations can initiate stable, periodic, long-wavelength patterns.

**Figure 3:**
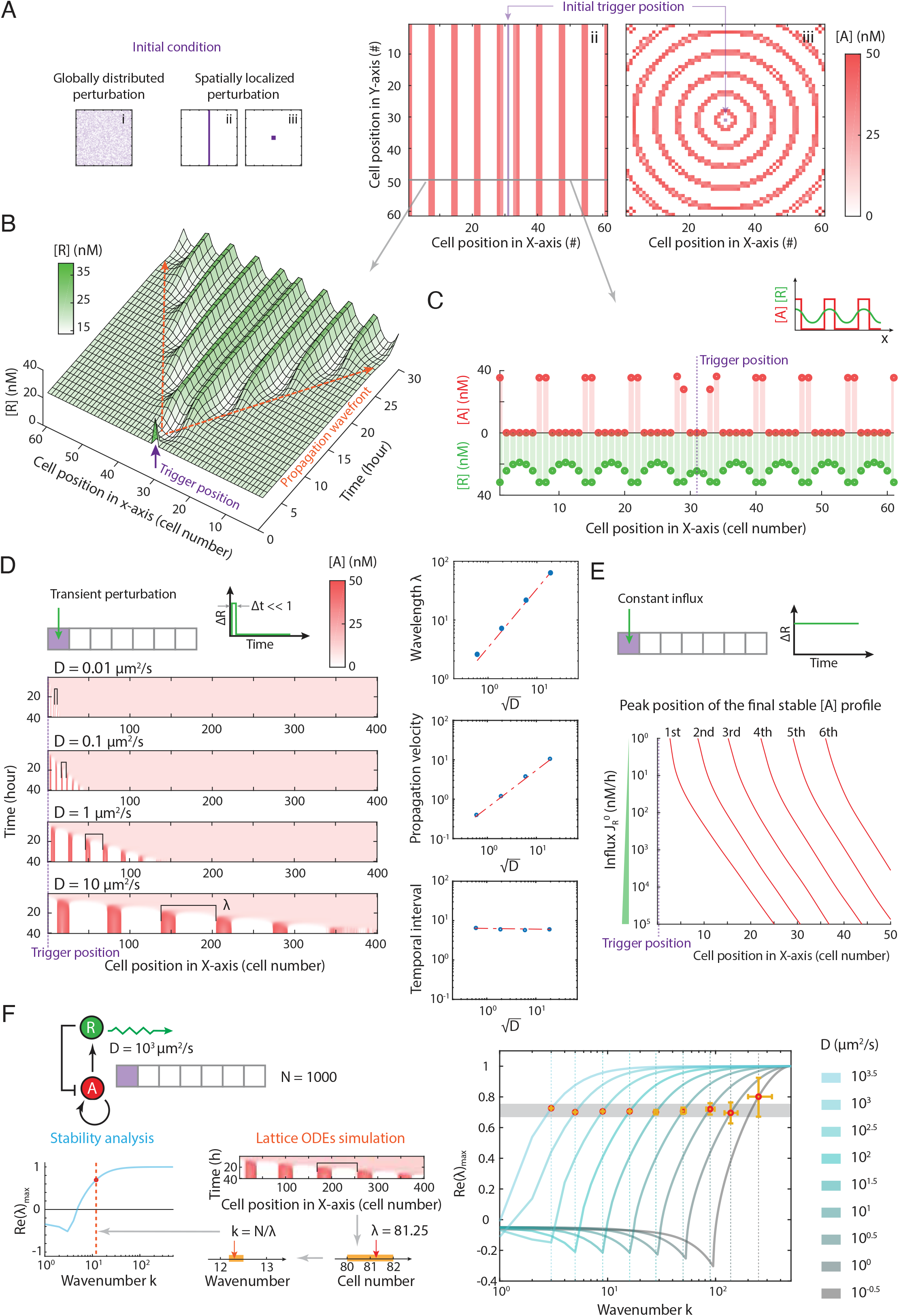
Spatially localized perturbation triggers low-wavenumber periodic patterns with the propagative dynamics. (A) **Spatially localized perturbation as initial condition provokes low-wavenumber regular patterns.** The global noise triggers synchronized high-wavenumber patterns. Meanwhile, the perturbing at one cell or a line of cells would promote stripes patterns and circle patterns, respectively. The final dominant wavenumber of stable patterns is notably lower than the ones from patterns triggered by global noise. (B) **This low-wavenumber pattern forms in a propagative fashion from the perturbation position.** A uniform speed traveling wavefront provokes a series of low-wavenumber stripes to form with uniform time intervals. The established patterns stay unchanged in both position and amplitude. (C) **The concentration of the activator and the repressor in the final pattern stay in phase.** The activator concentration profile behaves close to a binary style, while the repressor profile is more smooth and moderate. (D) **The diffusion coefficient determines the pattern wavelength.** A lower diffusion coefficient produces shorter wavelength patterns. The pattern wavelength measured from simulations with various diffusion coefficients is proportional to the square root of the corresponding diffusion coefficient. (E) **The intensity of the constant repressor influx alters the phase of pattern stripes.** The constant influx of repressors at the trigger position also induces propagative pattern formation. Increasing the influx level enlarges the distance from the trigger point to the first stripe. Meanwhile, the wavelength stays consistent. (F) **The dominated wavenumber measured from the final pattern is consistently lower than the wavenumber preference from stability analysis.** The evaluation repeated with various D indicates the actual patterning dynamics do not choose the one with the highest tendency but the one with 70% of the maximum, which locates at the edge of the plateau reading from the dispersion relation curve. The orange region shows the error bar from the measurement.

One of the most important features of patterns is their spatial periodicity. In our simulations, the wavelength of the single-morphogen pattern exhibited a square-root dependence on the diffusion coefficient, D, with larger diffusion coefficients producing larger intervals between adjacent stripes (Figure 3D). This relationship can be understood from a dimensional argument: D, with units of μm^2^/s, is the only parameter in the system whose dimensions include length. The velocity of pattern propagation is proportional to the square root of D for the same reason (Figure 3D). By contrast, the temporal interval between the formation of adjacent stripes is independent of D (Figure 3D).

In a biological context, the pattern-establishing perturbations could be generated by a group of “source” cells that produce the morphogen constitutively, as occurs with the sonic hedgehog morphogen in the developing neural tube (Dessaud, McMahon, and Briscoe 2008; Alaynick, Jessell, and Pfaff 2011; Briscoe and Small 2015). To understand how these source cells impact patterning, we simulated the system with a constant influx of morphogen at a defined position (Figure 3E). Higher morphogen levels suppress the expression of A in correspondingly larger regions around the source, effectively pushing out the position at which periodic patterning begins (Figure 3E). When morphogen influx is stopped after patterning is established, the existing patterned region expands inward, effectively “filling in” the proximal spatial region without affecting the established phase (Figure S3). Thus, morphogen production rates from source cells determine the phase of the final pattern, without affecting its period (Figure 3E). Taken together, these results indicate that a single-morphogen reaction-diffusion system, triggered by a spatially localized perturbation, can generate stable periodic patterns whose wavelength and phase can be controlled.

More generally, from the point of view of mode selection (Cross and Hohenberg 1993), these dynamics suggest that transient perturbations can selectively amplify modes with intermediate wavenumbers despite the monotonically increasing dispersion relation, as shown in a simplified one-dimensional example (Figure 3F). This contrasts with the classic stationary-pattern formation scenario, in which the dispersion curve is only positive for intermediate wavenumbers (Figure 1B left, purple lines). In that case, the final wavenumber of the emerging pattern results from the competition between the linear instability, the nonlinear terms, and the boundary conditions (Dee and Langer 1983; Murray 2001; Ben-Jacob et al. 1985). In our case, simulations revealed that the wavenumber of the spatial pattern corresponded to a Lyupanov exponent of about 70% of its maximum value (Figure 3F, right panel, gray shaded line).

### Spontaneous single-morphogen pattern formation is sensitive to noise

We next asked if single-morphogen pattern formation is sensitive to stochastic fluctuations (noise) in circuit components (Eldar and Elowitz 2010; Kilfoil, Lasko, and Abouheif 2009) (Figure 4A). To quantify this sensitivity and understand how it impacts patterning, we considered a version of the circuit that could be implemented synthetically, in which A and R are co-expressed from a single promoter (e.g. using an internal ribosome entry site (IRES) or 2A ribosomal skip sequence). In this configuration, the expression of the two genes can fluctuate due to noise, in a largely correlated manner due to their co-transcriptional expression. We therefore assumed that the dominant source of the noise was extrinsic, comprising both static variation as well as dynamic, correlated variability in the expression of the two genes (Thattai and van Oudenaarden 2002; Raser and O’Shea 2005; Elowitz et al. 2002). More specifically, we modeled expression noise using an Ornstein–Uhlenbeck process (Gillespie 1996; Rohlfs, Harrigan, and Nielsen 2013; Fox et al. 1988), which describes fluctuations with a standard deviation over time of σ, a standard deviation of mean expression across cells of η, and an autocorrelation time for fluctuations of *τ* (Figure 4B). These fluctuations are encapsulated in the random function H(i,t), where *i* denotes the cell index, and *t* denotes time, to produce a modified set of circuit equations:

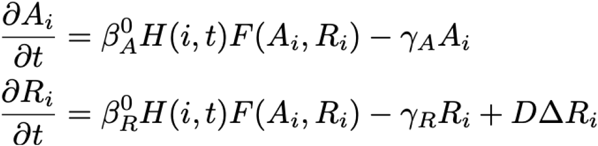

Here, H(i,t), the random time series for an individual cell, exhibits a normal distribution with mean m_i_ = <H(i,t)>_t_ and variance σ^2^. Mean expression levels are assumed to be normally distributed with mean 1 and variance η^2^. The timescale of fluctuations, *τ*, is taken to be about 10 hours, comparable to the mRNA half-life, which sets a natural timescale for fluctuations.

**Figure 4:**
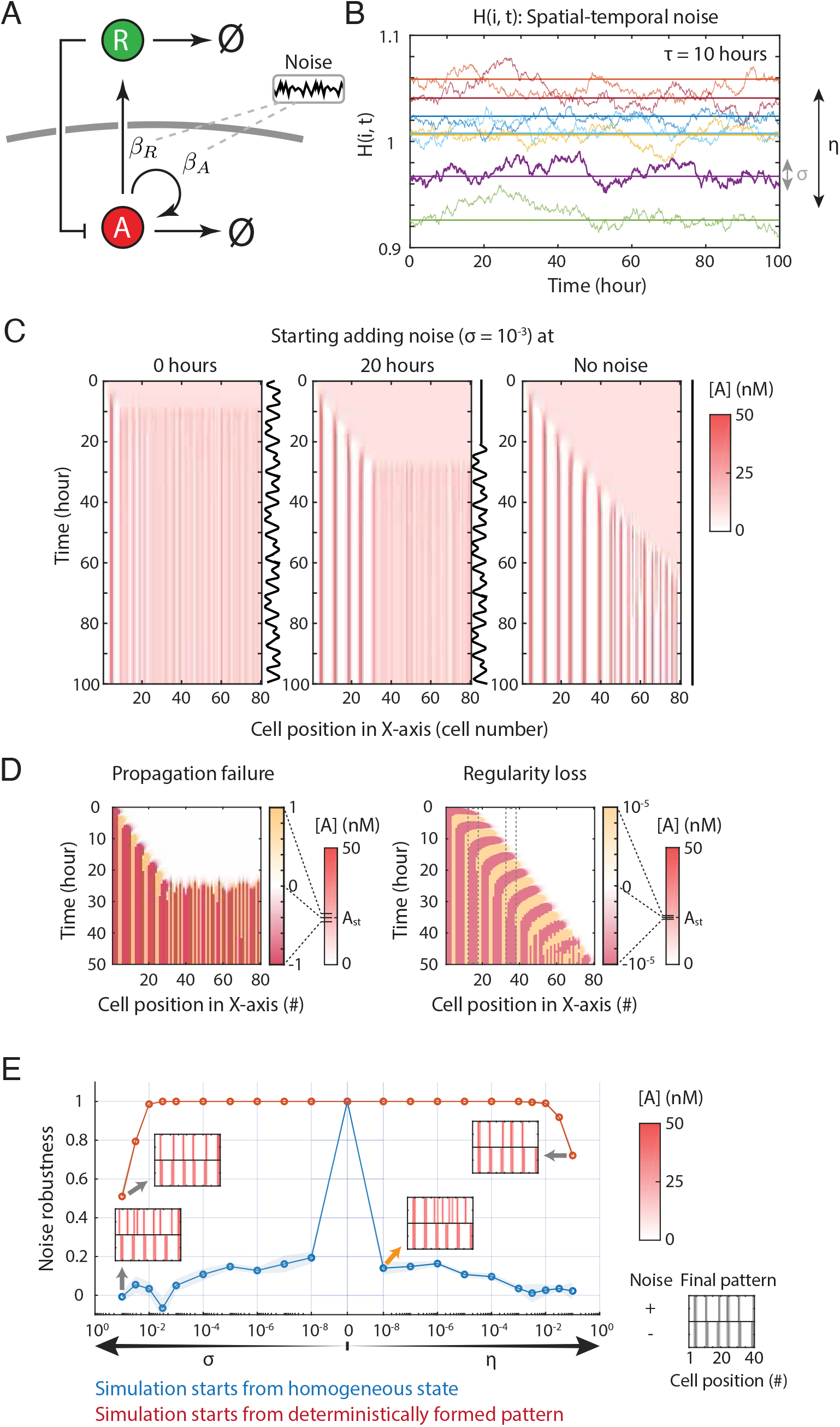
The propagation pattern is sensitive to noise during the formation process and is robust to noise once the pattern is established. (A) **The simulation of pattern formation in noisy conditions introduces noise in the production rate of both species**. The noise in the simulation recapitulates the fluctuation in protein production in general. (B) **The dynamic noise represents the temporal fluctuation in a particular cell. Meanwhile, the static noise describes the heterogeneity among the cell population.** The standard deviation quantifies the noise intensity, denoting σ and η, for the temporal dynamic noise and the static heterogeneity, respectively. (C) **The final pattern is sensitive to the timing of introducing noise.** The simulations take place on a 1×100 lattice with reflective boundary conditions and transient perturbation on the left end. At time 0 hours or 20 hours, σ = 0.001 level of noise starts to add to the simulation. The kymograph shows the pattern formation dynamics averaging from 20 repeated simulations. (D) **The noise triggers cells away from the perturbation center to undergo unwanted bifurcation, developing salt and pepper patterns and causing propagation failure. Meanwhile, the sequential activation eventually loses the regularity because the initiation trigger generates slightly different activation dynamics.** (E) **The minimal SMRD circuit is sensitive to noise only during pattern formation.** The spatial correlation of two final patterns with and without noise reveals the influence of noise and further indicates the circuit robustness to the noise. The simulations starting from the homogeneous state capture the pattern forming dynamics. Meanwhile, simulations with an established pattern as the initial condition reveal the pattern sustaining stage. During the patterning, the robustness score drops vastly even with a trace level of noise (orange arrow). The established pattern shows high robustness even with intensive noise.

In the presence of the noise sources described above, our simulations show that even small amplitudes of noise disrupted pattern formation (Figure 4C). In a one-dimensional system with stochastic fluctuations (σ = 0.001, η = 0), patterning was disrupted after formation of the first stripe (Figure 4C, left vs. right panel). Suspending noise until the middle of the simulation (t = 20 h) allowed more stripes to form, but pattern propagation was quickly blocked once the noise was present (Figure 4C, middle panel). On the other hand, while noise prevented the formation of new stripes, it did not disrupt stripes that had already formed. These results indicate that noise can disrupt pattern formation and cause propagation failure without destabilizing patterned regions (Figure 4D, left). Meanwhile, we also noticed that under the noise-free condition, the pattern eventually loses its regularity (Figure 4C, right). The distal cells experience different activation dynamics compared with proximal cells. This “initiation effect” perturbs the spatial-temporal periodic cycles, causing regularity loss in distal regions.

To quantify the ability of noise to disrupt pattern formation, we analyzed patterning across different values of σ and η. As a measure of noise robustness in pattern formation, we analyzed the spatial correlation between the steady state patterns with and without noise. Values near 1 indicate patterning is unaffected by noise, while values near 0 indicate the absence of periodic patterning at the expected frequency. With a homogeneous initial condition, the circuit was sensitive to noise during pattern formation (Figure 4E, blue line). Disruption increased with the amplitude of noise, but even extremely low values of σ or η were sufficient to at least partially disrupt patterning (Figure 4E, orange arrows). By contrast, if we initialized the system with a deterministically formed pattern, even larger values of noise, up to at least ∼10%, failed to disrupt it (Figure 4E, red line). These results were again consistent with the notion that noise disrupts patterning specifically in unpatterned regions and, more generally, could be understood in terms of the ability of noise-stimulated patterns to fill in previously unpatterned regions, but not penetrate regions in which the pattern is already formed (Figure 4C, middle panel).

### An inhibition-release mechanism overcomes noise sensitivity

Based on the results above, and inspired by aspects of retinal patterning in *Drosophila (Heberlein, Wolff, and Rubin 1993a),* we reasoned that suppressing A and R expression in unpatterned regions could prevent their premature activation by noise, while preserving their ability to pattern in response to the advancing wave of pattern formation. To implement this behavior, we added an “inhibition release” mechanism to the A-R circuit, based on a hysteretic positive feedback loop (Figure 5A). More specifically, we incorporated an intracellular regulator, denoted M, which can exist in either an inactive (M) or active (M*) state. We assume three key interactions for M. First, the transition from the inactive M to active M* is stimulated by R in an ultrasensitive manner. Second, M* stimulates M activation, creating a positive, ultrasensitive feedback loop that exhibits bistability. Third, M* inhibits A degradation, which is assumed to occur at an accelerated rate, and thereby boosts A protein levels. With these assumptions, in the appropriate parameter regimes, M* exhibits an ultrasensitive, hysteretic dependence on its input, R (Figure 5A). Low levels of M* lead to correspondingly low levels of A, which in turn ensures low levels of R production, maintaining the cell in a ground (pre-patterned) state. By contrast, when M* is high, stabilization of A allows the circuit to exhibit the pattern forming behaviors described above (Figure 5B, phase diagram). In this way, an advancing wave of R morphogen can convert M to its active form, progressively de-inhibiting the A-R circuit as it propagates (Figure 5C).

**Figure 5:**
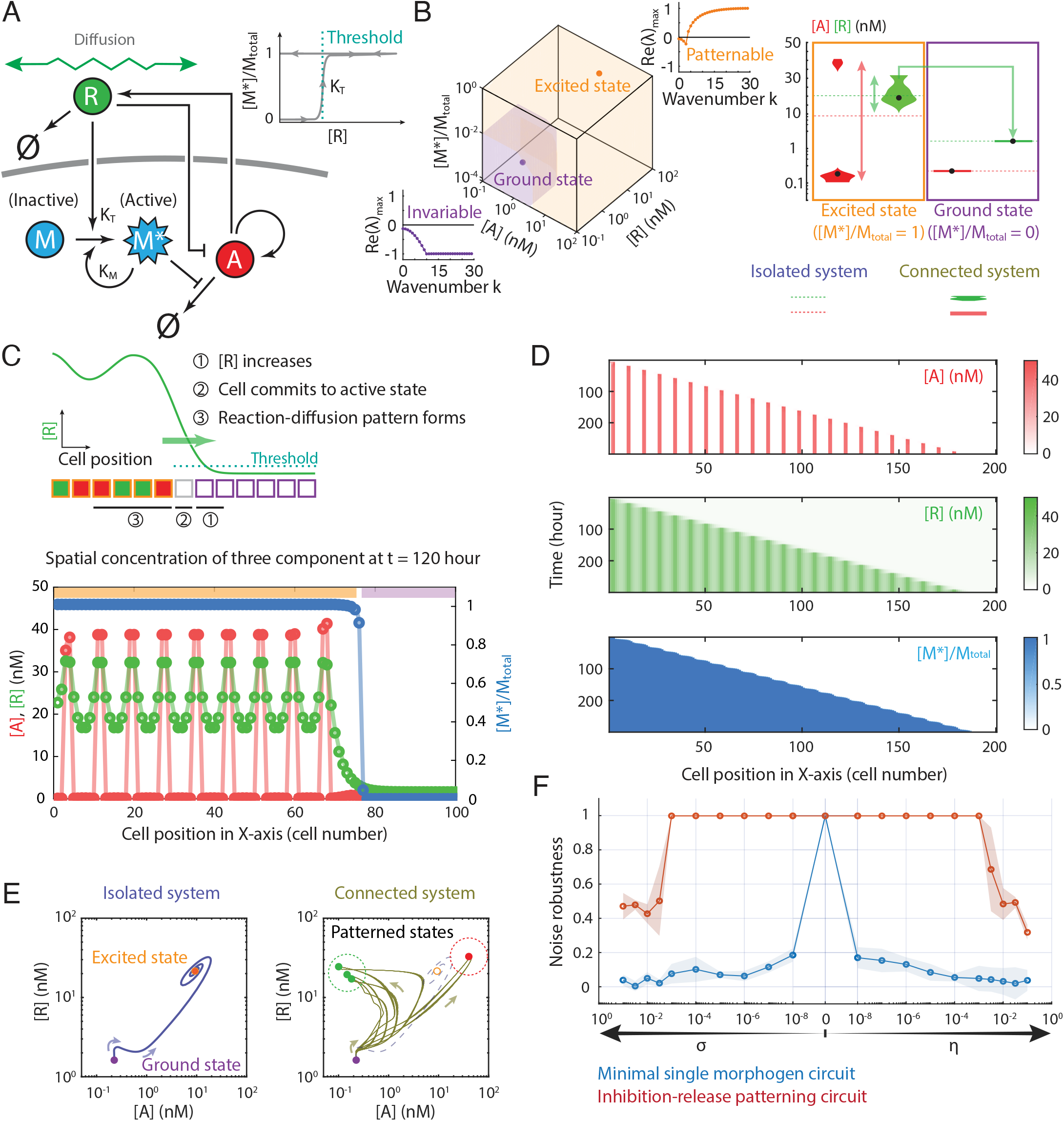
A bistable switch transiently stabilizing the patterning cells prevents the noise-triggered irregular bifurcation dynamics. (A) **Implementing this inhibition-release mechanism, we propose a three-components circuit that relies on an intracellular regulator (M) to temporarily stabilize pre-patterning cells.** The third component has two forms: M* (active) and M (inactive). Only the active form could repress the degradation of A and restore its level in the cell. Meanwhile, both R and M* mediate the transition from M to M*. The ultra-sensitivity of R-mediated M activation ensures the M* level is either low or high. (B) **This inhibition-release circuit is bistable in the isolated system (no diffusion). The excited state is patternable while the ground state is always stable in the connected system.** With proper parameters, the switch between two forms of M is designed to accomplish either stabilizing pre-patterning cells (ground state) or releasing the inhibition to form propagative periodic patterns (excited state). Both states are stable with homogeneous A and R levels (dash lines) in the isolated system. Meanwhile, unlike the ground state cells staying stable, the excited state cells bifurcate forming two clusters in the connected system(solid lines or shapes). **(C) The patterned cells convert adjacent ground state cells and trigger new stripes in a propagative fashion.** Initially, all cells stay in the ground state. The local increase of R converts the cell to the excited state. Reaction-diffusion pattern forms on these patternable cells. The high concentration of R from the region of patterned cells diffuses to their ground state neighbors and further converts them to be patternable. Three events, the repressor elevation in the ground state, the transition of the cell state from ground to excited, and the reaction-diffusion pattern formation on newly excited cells, occur in a synchronous sequential fashion. A snapshot of the concentration profiles from the simulation captures the detailed dynamics of the pattern formation, confirming our proposed dynamics in the model design. (D) **The inhibition-release circuit generates propagative periodic patterns, validated by the lattice ODE simulation.** The simulation kymograph presents similar dynamics of A and R to the minimal SMRD circuit, where the periodic stripe pattern forms sequentially following the constantly propagated wavefront. Meanwhile, cells transform from the dormant to the active state, ahead of the wavefront swapping and patterning. (E) **During patterning in the connected system, cells bifurcate after escaping from the ground state while bypassing the excited state.** The phase portrait of the connected lattice system shows the populational behavior of bifurcation. Instead of ending at a single stable steady-state as in an isolated system, once cells escape from the ground state, they end up with multiple stable states. All final states fall into two clusters, responding to a typical periodic pattern’s bright (red dash circle) and dark regions (green dash circle). (F) **The inhibition-release circuit vastly increases the patterning robustness to either type of noise.** Pattern formation simulations under such a paradigm at various noise intensities show better robustness to either type of noise than the minimal circuit. For minimal circuits, the regularity of the pattern is sensitive to even considerably low-level noise. Meanwhile, the inhibition-release circuit shows robustness to a wide range of both types of noise.

To test whether this circuit could prevent noise from triggering premature circuit activation, while still allowing pattern propagation, we simulated the system on a one-dimensional cell lattice (Figure 5C, D). We initialized the lattice with all cells in the inhibited ground state (low M*), which remained stable across the lattice in the absence of external R (Figure 5B, purple box). Production of R at the boundary triggered activation of M*, and consequent de-inhibition of the A-R circuit, in boundary-adjacent cells (Figure 5C). Since excited state cells produce R at a higher rate than ground state cells (Figure 5B, right panel, orange box), the R concentration gradient drove a net R flux towards more distal, inhibited cells, leading to a propagating front of M* activation (Figure 5C, 5D). Once de-inhibited in this way, the A-R circuit progressively developed periodic patterns of A and R concentration through a diffusion-driven bifurcation. Patterning occurred through three sequential stages: (1) elevation of the repressor, R, in the inhibited ground state, prior to its activation; (2) transition of the cell from the ground to the excited state by activation of M, and (3) reaction-diffusion pattern formation by the A-R circuit in the excited cells (Figure 5C). The third step resembled periodic stripe patterns following a propagating wavefront in the minimal A-R circuit (Figure 3). The effects of R diffusion can also be observed in the phase portrait (Figure 5E). Once cells escape from the inactive state, instead of ending at a stable excited state as in an isolated system, they exhibit multiple stable states as a consequence of the diffusion-driven bifurcation. Most importantly, all final states fall into two clusters, responding to the typical bright and dark regions of a periodic pattern (Figure 5C).

The inhibition-release mechanism makes patterning more robust to noise. We simulated the circuits with or without inhibition release at different noise intensities. Without inhibition release, even very low levels of noise were sufficient to disrupt patterning, as discussed previously (Figure 5F, blue line). With inhibition release, patterning remained robust across many orders of magnitude of noise strength that disrupted the simpler circuit (Figure 5F, red line). Taken together, these results show that single-morphogen patterning can overcome the challenge of noise-induced spontaneous high-frequency patterns.

### Growth-coupled patterning is robust to noise

We explored in what follows another mechanism through which single-morphogen reaction-diffusion patterns can be robust to noise, that relies on tissue growth. In many developmental systems, such as somitogenesis or intestinal crypt development, tissue patterning is closely coupled to growth (Schnell et al. 2002; Moreno and Kintner 2004; Itzkovitz et al. 2012). In some of these systems, including intestinal crypts and apical meristems, progenitor cells at one location undergo repeated asymmetric divisions generating the rest of the tissue, which patterns as it is produced. We reasoned that similar growth-coupled patterning could reduce the noise susceptibility of single-morphogen patterning. Assuming tissue growth occurs at a similar or slower rate compared to pattern propagation, newly formed tissue can immediately be patterned by adjacent, already patterned, tissue regions. In such a dynamically growing tissue context, the single-morphogen circuit could operate in a regime in which cells pattern immediately after division, and do not “wait” in a noise-sensitive state for a propagating front of patterned cells to reach them (Figure 6A).

**Figure 6:**
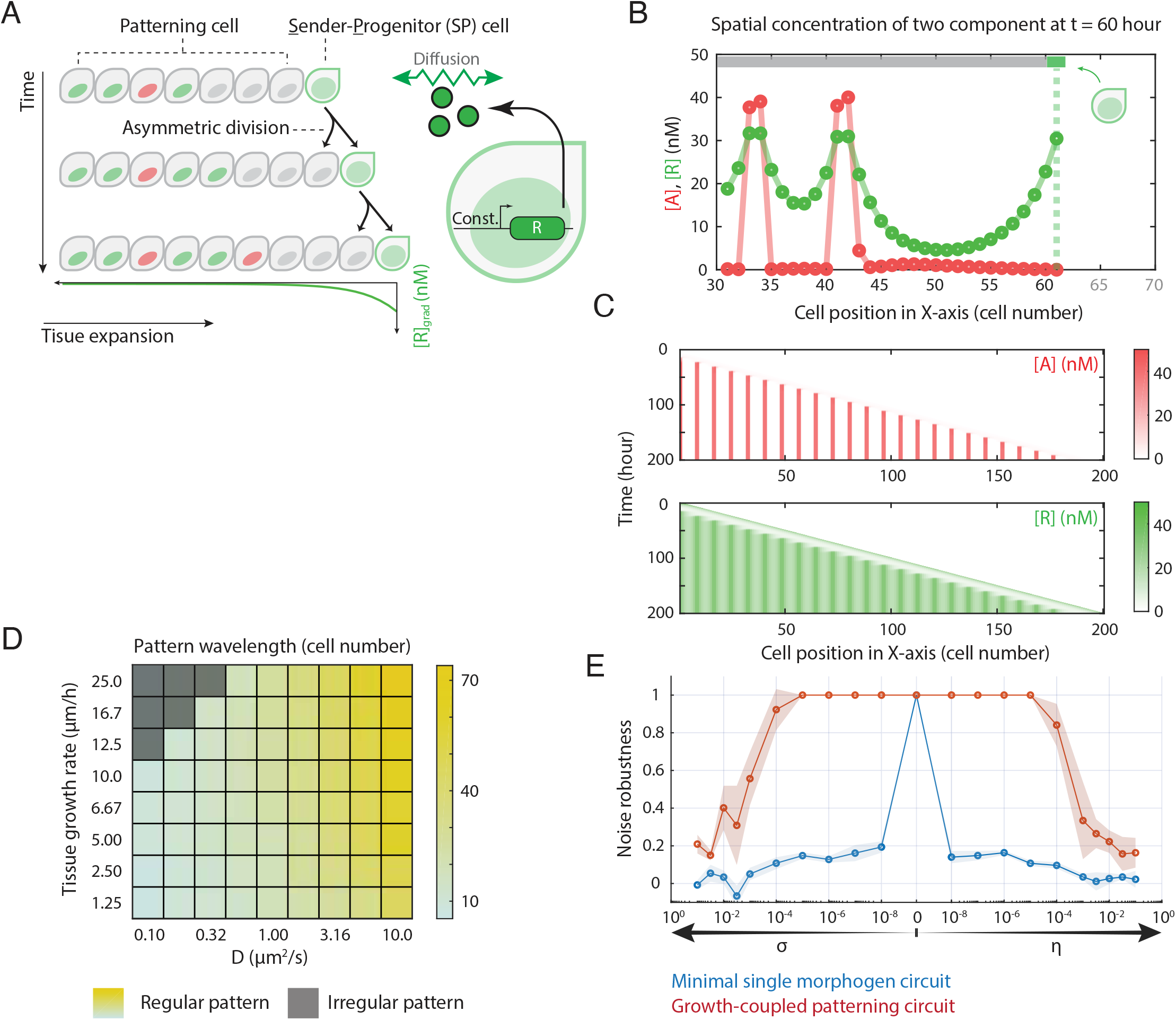
Coupling the tissue growth with the SMRD circuit increases the noise tolerance of pattern formation. (A) **The growth-coupled patterning model assumes a tissue growing in one dimension.** All patterning cells with the minimal single-morphogen patterning circuit originate from a specialized cell type, the sender-progenitor(SP) cell, which can asymmetrically divide, creating a patterning cell while keeping its number constant. The newborn cell appears between the SP and the elder patterning cell so that the entire tissue elongates as the consequence of cell division. Meanwhile, the SP cell stays at the tissue’s elongation end and produces the repressor constantly. The Repressor gradient established by the SP cell stabilizes newborn patterning cells. [R]_grad_ is the spatial concentration profile of the repressor gradient from the SP cell as an exponential decay function. (B) **The growth-coupled circuit forms periodic patterns on the actively growing tissue.** The pattern formation process, initiated by the SP cell, takes place on the patterning cells. The new stripes emerge periodically near the growing center, where new patterning cells are produced. Meanwhile, the stabilized patterns on the mature pattering cells stay unchanged. (C) **The growth-coupled circuit generates periodic patterns on expanding tissue, validated by the lattice ODE simulation.** The simulated kymograph presents similar dynamics of A and R to the minimal single-morphogen patterning circuit. The propagation speed of pattern stripes is coupled with tissue growth rate. (D) **Both diffusion coefficient and tissue expansion rate scale the pattern wavelength.** Besides the diffusion coefficient, the tissue expansion rate is the second spatial parameter in the model. A high expansion rate widens the wavelength. The diffusion coefficient is well-coupled with a wide range of tissue growth rates. Patterning circuits with small D fail to form regular stripes patterns on a fast-growing tissue. (E) **The growth-coupled patterning circuit notably increases the noise robustness of the single-morphogen patterning.** The noise interference test repeated with the tissue-growing paradigm shows significant improvement of noise robustness during the pattern formation stage. When the noise is beyond the tolerance, the robustness score drops faster for extrinsic noise than intrinsic noise.

To test whether tissue growth could allow robust single-morphogen patterning, we analyzed the minimal A-R patterning circuit on a growing cell lattice. We designated a specialized sender-progenitor (SP) cell type that asymmetrically divides to generate a differentiated cell, which inserts between the SP and the products of the previous asymmetric divisions (Figure 6A). This simple toy model of tissue growth maintains a constant number of SP cells at one end of the tissue, while generating a linearly elongating lattice of patterning-competent cells. We assume that only the differentiated cells express the full circuit, and are therefore competent to form patterns. Finally, in addition to its role in producing the tissue, the SP cell also serves as a constitutive producer of the R repressor, establishing a concentration gradient whose length scale extends several cell diameters into the most recently generated cells (Figure 6A). In this scheme, the extracellular repressor emanating from the SP cell inhibits patterning in the proximal region (Figure 6B). However, as tissue growth progresses, individual cells eventually emerge from this inhibitory zone and can begin to pattern. Together, these features could allow growth-coupled patterning and avoid the disruptive effects of noise in pre-patterned cells.

To simulate deterministic growth-coupled patterning, we initialized the cell lattice with a single SP cell, and then periodically inserted newly differentiated cells directly adjacent to it (Figure 6B). In the differentiated cells, we simulated the same A-R circuit with the same parameter set analyzed above (Figures 3,4), imposing reflective boundary conditions at both ends of the growing tissue. As anticipated, newborn cells initially expressed low levels of both A and R due to the production of R by the adjacent SP. Once a cell reached a distance from the SP, its A and R production rates switched to either a high state, in which both proteins were strongly expressed, or a low state, in which both were weakly expressed, depending on the states of the previous adjacent cells (Figure 6B). This led to periodic patterning (Figure 6B, 6C). Patterning proceeded indefinitely, and no defects were observed in this deterministic case for these parameters (Figure 6C). Further, growth-coupled patterning occurred across a broad range of values of diffusion coefficients and tissue growth rates (Figure 6D). These results show that coupling to growth can allow progressive patterning.

We next asked whether growth-coupled patterning could overcome the disruptive effects of noise. We simulated the system with the same growth dynamics, but systematically varied the levels of temporal and static noise, as discussed previously. Across a wide range of noise intensities, the growth-coupled system maintained predictable patterning for conditions that prevented periodic patterning in static lattices (Figure 6E). These results show that coupling to tissue growth can allow predictable patterning with a single morphogen in the presence of noise.

## Discussion

Turing patterning is a classic concept in developmental biology. In this work, we investigated Turing patterning using only a single diffusible morphogen — a regime that does not permit patterning under standard assumptions. Implemented on a discrete cell lattice, rather than in a continuous medium, this single-morphogen Turing system can spontaneously generate patterns ranging from spatially alternating cell states to irregular, longer wavelength features (Figure 2). Motivated by the powerful roles that initial conditions and boundary conditions play in development (Hiscock and Megason 2015; Murray 2001), we considered their impact on patterning by this circuit. When stimulated by a point or line initial condition, the same circuit can generate propagating “stripes” that, once patterned, remain stable even if the initiating signal is removed (Figure 3). Transient initial conditions can thus control what types of single-morphogen circuits are capable of patterning.

In the single-morphogen system, low spatial frequency patterning modes must compete with the global attractor of high spatial frequency noise. Without mitigation, noise dominates, generally limiting the range and precision of the low frequency patterns. More specifically, we found that noise was most disruptive when it had time to trigger high spatial frequency patterns that block the propagation pattern from the initiating boundary (Figure 4C). However, two remarkably simple, biologically plausible mechanisms are sufficient to ensure robust patterning.

On a static cell lattice, cells could circumvent noise-induced high-frequency patterns by “locking down” circuit activity until the propagating front of pattern formation approaches sufficiently to provide a high enough local concentration of morphogen to “unlock” the circuit. At that point, the adjacent cells are sufficiently well patterned to ensure the correct outcome for the cell in question. While this effect could be achieved in many ways, here we demonstrate how the a relatively simple bistable switch, comprising a single positive autoregulatory factor, can provide this functionality, ensuring robust patterning (Figure 5).

In growing tissues, noise can be circumvented in a distinct manner, by limiting patterning with tissue growth (Figure 6). In this regime, a set of progenitor cells elongate the tissue by self-renewing and differentiating into patterning cells. These newborn patterning cells immediately respond to the local environment to reach the appropriate state. In this case, cells spend little time in a noise-susceptible state where they are patternable, but not yet patterned. Hence, such a regime avoids the unwanted formation of high-frequency patterns.

How realistic are these assumptions? Mathematically, multicellular patterning systems are often imagined to first approach a transient homogeneous unstable state before cells diverge into distinct fates (Sprinzak et al. 2011; Raspopovic et al. 2014; Onimaru et al. 2016; Palau-Ortin et al. 2015). However, in natural development, patterning may more often proceed from a pre-patterned state that avoids the need for further symmetry breaking. In fact, many biological systems exhibit features that appear similar to those considered here. For example, in drosophila retinal development, the periodic patterning of ommatidia occurs dynamically in the wake of the morphogenetic furrow, as it sweeps across the eye disc, from posterior to anterior, over two days (Heberlein, Wolff, and Rubin 1993b). Cells located anterior to the morphogenetic furrow are unpatterned, while cells posterior to the morphogenetic furrow assemble into patterned ommatidia in a stepwise process. The progressive unlocking of patterning by the furrow is analogous to our inhibition-release model. We note, however, that this system relies on both the hedgehog and Dpp signaling pathways, rather than a single-morphogen, as analyzed here.

Patterning of the intestinal crypt provides an example of growth-coupled patterning (Gehart and Clevers 2019). In this case, the columnar cells at the crypt base act as progenitor cells, dividing asymmetrically to produce a continually growing crypt with patterning of differentiated absorptive (high Notch signaling) or secretory (low Notch signaling) cells. Here, growth appears to play a role in unlocking competence for patterning. Further, Wnt ligands secreted by Paneth cells at the base of the crypt play a key morphogenetic role in inhibiting differentiation, although other diffusible signals also play roles in this system.

Finally, in plant development, apical meristem development involves a central zone of progenitor cells that divide and differentiate to replace cells in the peripheral and rib zones, which divide more rapidly and further differentiate (Nakajima and Benfey 2002; Carles and Fletcher 2003). Meanwhile, the shoot apical meristem also produces the signaling molecule auxin, preventing axillary buds, which initially locate in the leaf axil dormant, from undergoing shoot development because of the apical dominance (Dun, Ferguson, and Beveridge 2006; Kebrom 2017).

The simplicity of single-morphogen patterning could make it ideal for the nascent field of synthetic development. As synthetic biology progresses from single-cell to multicellular systems, a key challenge is engineering cells that can establish their own spatial patterns. In this context, circuit architectures with minimal components and interactions, such as the single-morphogen patterning systems explored here, are critical. The entire single-morphogen patterning circuit proposed here could be implemented with well-characterized components. For example, Gal4 or engineered zinc finger transcription factors (Zhu et al. 2021; Khalil et al. 2012) could serve as the activator, A. The repressive morphogen, R, could be implemented with the combination of a secreted diffusible signaling molecule and an intracellular repressor whose expression it activates. Sonic Hedgehog has been shown to form well-defined morphogenetic gradients in 3T3 cells, making it a candidate for the morphogen (Li et al. 2018). Other natural pathways such as BMP could also function in this way. Alternatively, synthetic small signaling molecules such as auxin (Ma et al. 2020; Liang, Ho, and Crabtree 2011), and variants of the synNotch system that allow morphogenetic patterning (Morsut et al. 2016) could enable the engineering of patterning systems orthogonal to natural pathways. Tet repressor variants could allow downstream negative regulation. To circumvent noise, the bistable switch (Figure 5) could be implemented with a self-stabilizing protease (Gao et al. 2018), synthetic phosphorylation systems (Mishra et al. 2021; Woodall et al. 2021), transcriptional autoregulation (Zhu et al. 2021), or RNA-level regulation (Levine et al. 2007; Morris and Mattick 2014; Dykes and Emanueli 2017), among other mechanisms. Finally, coupling patterning to tissue growth could be realized using spatially confined cell culture systems that restrict growth to elongated channels, as was shown for mammalian cells using the “mother machine” microfluidic system (Pearl Mizrahi et al. 2016; Potvin-Trottier et al. 2016). Thus, we anticipate this circuit providing a feasible foundation for engineering synthetic pattern formation systems. Turing’s remarkably general pattern-forming architecture is well known for the broad range of patterns it can generate. It is interesting to see how initial conditions, noise, and variations in circuit architecture jointly allow robust patterning in both natural and synthetic multicellular contexts.

## Methods

### Lattice ODE simulation

#### Framework

To tie experiment and modeling closely, we use cell-lattice simulation to approximate experimental conditions. Depending on the geometry of the simulation, we run simulations on 1-D or 2-D. In the one-dimensional case, virtual cells in a finite length chain are connected with their left and right neighbors. In a two-dimensional case, cells are connected in a square lattice. For each cell, we assign one set of ODEs. Meanwhile, morphogen diffusion is realized by molecules transferring among adjacent cells.

#### Equilibrium simulation

An initial concentration profile guess is assigned to all components. In the pre-equilibrium simulation, diffusion is turned off and the ODE systems in each virtual cell would evolve by themselves. The system eventually reaches the homogeneous steady state, in which all the cells have the same concentration profile in each circuit component.

#### Patterning simulation

To start the pattern formation, we turn on the diffusion. A specific trigger, corresponding to the initial condition, creates a transient heterogeneous perturbation in the system. The simulation finishes once the pattern stops changing spatially and temporally.

#### Parameter estimation

To better fit the biological context, we estimate model parameters from BioNumbers (Milo et al. 2009) and from our own wet-lab experiments. The estimated parameter values are shown in Table 1. Our model assumes that the cell size is 10 μm x 10 μm x 10 μm, and we further deduce other parameter values from there.

**Table 1:** Parameter list and estimated values in simulations

| Parameter Name | Parameter Description | Parameter value |
| --- | --- | --- |
| $\beta_A$ | Production rate of activator | 100 nM/h |
| $\beta_R$ | Production rate of repressor | 80 nM/h |
| $\gamma_A$ | Degradation rate of activator | 1.5 /h |
| $\gamma_R$ | Degradation rate of repressor | 0.5 /h |
| D | Diffusion coefficient of repressor | 1 $\mu\text{m}^2/\text{s}$ |
| $K_A$ | Dissociation constant of activator | 10 nM |
| $K_R$ | Dissociation constant of repressor | 10 nM |
| K | Basal leakage of promoter | 0.01 |
| h | Hill coefficient of promoter | 2 |

#### Initial condition

Our simulations use two types of initial conditions, which lead to two types of patterning results and dynamics. All lattice nodes are subject to global random noise in the concentration of A (or R). The perturbation profile is generated by random sampling from a Gaussian distribution with zero mean (μ = 0) and given standard deviation (σ = σ_0_). The spatial localized initial condition only affects selected cells in the same fashion. The selected cells usually follow a certain geometry, either a single dot or a straight line. The pattern can be triggered by acting upon any of the two species, by increasing or decreasing its concentration for an arbitrary time window. Specifically, some tests involve a constant influx from a line of cells as the initial condition.

#### Boundary condition

We consider a square lattice of size *l × w* for our simulations (in some cases the system is one-dimensional by setting *w* = 1). The simulation is repeated with two types of boundary conditions: reflective and periodic. The general features of the SMRD pattern are not affected by the choice of boundary condition.

#### Noise interpretation

Our simulations consider two types of noise: temporal dynamic noise or static heterogeneity. Specifically, we generate the noise profile H(i,t)|_η, σ_ with given σ and η ahead of the simulation. The temporal dynamic noise is denoted by H(i,t)|_η = 0, σ_ and the static heterogeneity by H(i,t)|_η, σ = 0_. Initially, the noise randomizes the production rate of both species. Linear interpolation between time points in H(i,t)|_η, σ_ is used when needed in the simulation. In the case where the noise starts to interfere with the simulation at a later time point, we reset the value of H(i,t)|_η, σ_ to 1 during the noise-free time period.

#### Modeling growing tissue

The lattice is initiated with two lines of cells. One line is formed by cells in which the reaction-diffusion system is active. The other line is formed by sender-progenitor (SP) cells, which constitutively produce the repressor. After a time period T_cyc_, a new line of patterning cells is added between the SP cells and the patterning cells. The initial morphogen concentrations in patterning cells are the same as in the SP cells. The tissue expansion rate is calculated as the cell size divided by T_cyc_.

### Data Analysis Techniques

#### Final stable pattern auto-correlation

2-D spatial auto-correlation, calculated by MATLAB function corr2() on the original 200 x 200 lattice simulation final frame, reveals lateral inhibition (Figure 2B, 2C). In the 1-D pattern, the auto-correlation function is calculated with the MATLAB function corr().

#### Pattern wavelength determination

To measure the pattern wavelength, the MATLAB peak detection function identifies the peak positions from the auto-correlation profile of the 1-D simulation final frame. The measured wavelength is the average distance of two adjacent peaks.

#### Determination of the pattern propagation speed

We locate the peak positions in the final pattern and then record the time when the activator in the corresponding cells crosses a threshold (10% above the steady-state). The time series defines the propagation period. The propagation speed is the spacing between two peaks divided by the propagation period.

#### Noise robustness quantification

To evaluate the noise robustness of a certain model, we run the pattern formation simulation with and without noise. The spatial correlation between the two final repressor profiles quantifies the similarity of the two patterns. If the correlation is near 1, we claim the model is robust to this particular level of noise. Otherwise, if the correlation is close to 0, the model is sensitive to noise. This test is repeated at various levels of noise (both temporal dynamic noise and static heterogeneity) in all three models, to evaluate the noise robustness of three circuits.

## Acknowledgments

MBE is a Howard Hughes Medical Institute Investigator. This research was supported by the Allen Discovery Center program under Award No. UWSC10142, a Paul G. Allen Frontiers Group advised program of the Paul G. Allen Family Foundation.

J.G.O. was supported by the Spanish Ministry of Science and Innovation and FEDER, under project PGC2018-101251-B-I00, by the “Maria de Maeztu” Programme for Units of Excellence in R&D (grant CEX2018-000792-M), and by the Generalitat de Catalunya (ICREA Academia programme).

We acknowledge insightful feedback from Elowitz lab members. The authors have declared that no competing interests exist.

## Author contributions

Conceptualization and Investigation: S.W., J.G-O, M.B.E.

Formal Analysis, Software, and Visualization: S.W.

Writing: S.W., J.G-O, M.B.E.

## Code reporting

Simulation code and data is accessible from CaltechDATA. Wang, S., García-Ojalvo, J., & Elowitz, M. B. (2022). Periodic spatial patterning with a single morphogen (Version 1.0) [Data set]. CaltechDATA. https://doi.org/10.22002/D1.20060

