## Supplementary figures and images for "Periodic spatial patterning with a single morphogen"

### Movie S1

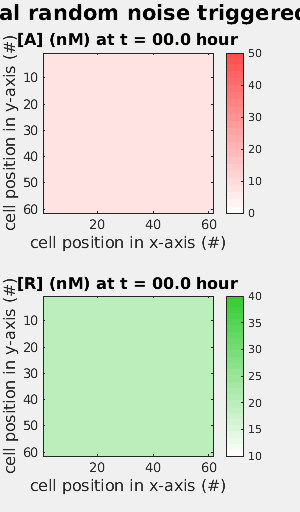

### Movie S2

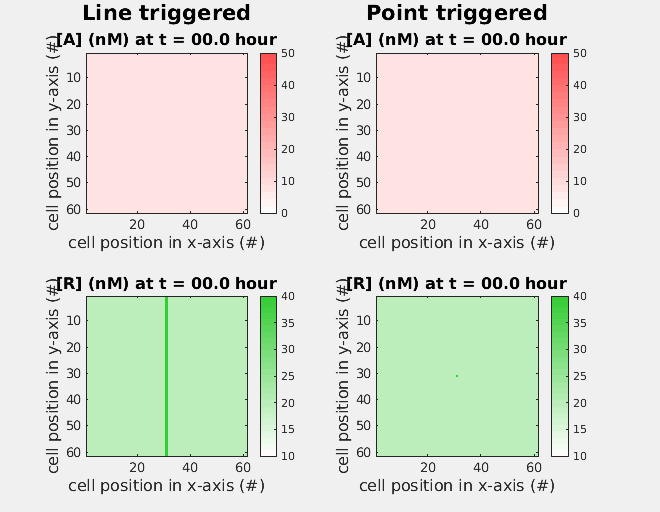
