## Appendix and Supplementary Figures for "Periodic spatial patterning with a single morphogen"

### Supporting information

S1 Appendix: Analytical analysis for single-morphogen patterning circuit in continuum

S2 Appendix: Stability analysis for single-morphogen patterning circuit in discrete lattice

S1 Figure: Single-morphogen patterning circuit with Gierer–Meinhardt kinetics

S2 Figure: Pattern initiation is robust with a wide range of perturbations

S3 Figure: Removal of morphogen influx causes pattern stripes “fill-in” the empty space

S1 Movie: Single-morphogen circuit patterning initiated with global random noise

S2 Movie: Propagation pattern formation triggered by localized perturbation

**S1 Appendix: Analytical analysis for single-morphogen patterning circuit in continuum**

The following analysis considers a two-component system, and the conclusion is valid for a multiple-component pattern. Given a single-morphogen patterning circuit,

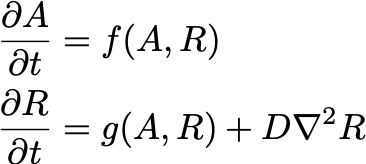

the necessary and sufficient condition to generate stable spatial patterns is that the Jacobian matrix stays stable without diffusion and becomes unstable for some wavenumber k with diffusion. denoting J(k) as

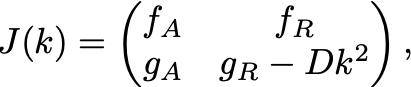

where k is the wavenumber (k = 1, 2, 3, …). The eigenvalue is the solution of

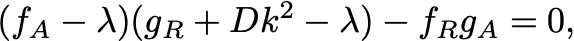

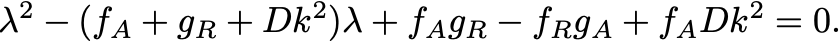

when k is large, the equation becomes,

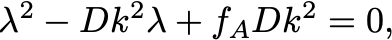

Assuming Re(λ_1_) ≤ Re(λ_2_), then

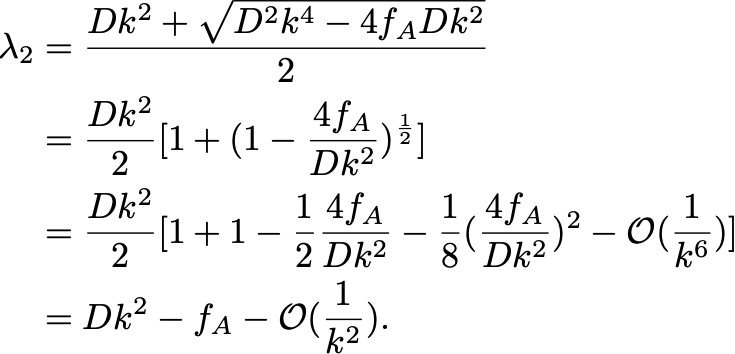

is monotonic increasing as the function of k when k is large.

**S2 Appendix: Stability analysis for single-morphogen patterning circuit in discrete lattice**

- **Steady-state of the isolated system**:

Given the lattice ODE of a single-morphogen patterning circuit, firstly we solve the steady-state without diffusion terms.

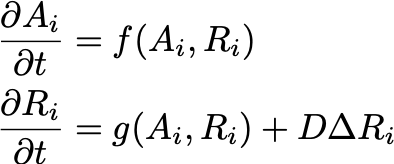

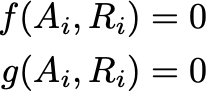

The steady-state concentrations, denoted as A_0_ and R_0_, are valid for all lattice nodes.

- **Stability analysis on 1-D or 2-D discrete lattice**:

Similar to the calculation in Plahte's work (Plahte 2001), denoting C = [A_1_, R_1_, A_2_, R_2_, ... A_N_, R_N_], the linearized kinetics of the whole system near the steady-state is

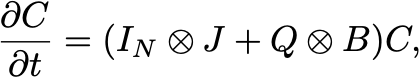

where

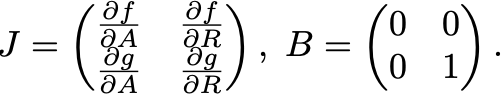

Q is the matrix describing the connectivity of the lattice. For all diagonal elements, Q_ii_ = - c, where c is the number of cells directly connected with cell i. If diffusive molecules can directly transfer between cell i and cell j, Q_ij_ = 1. Otherwise, Q_ij_ = 0. If the connectivity matrix Q can be diagonalized with matrix U and U^-1^, then we have,

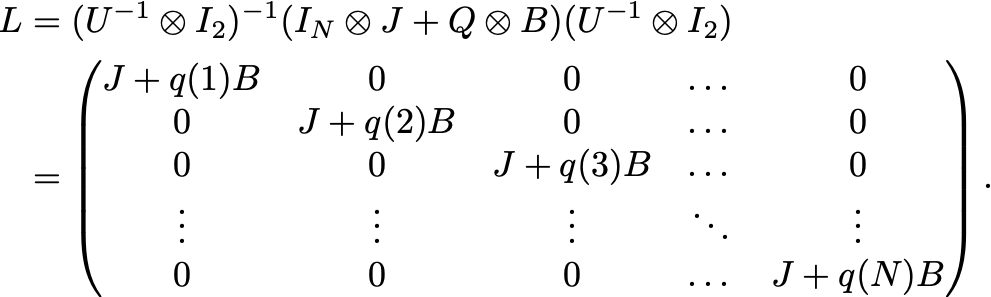

Since L is a block-diagonal matrix, which means,

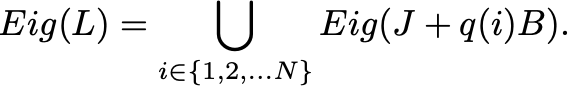

The whole lattice is stable, if and only if all of J+q(k)B are stable. Meanwhile, The lattice would be unstable as long as one of the J+q(k)B is unstable.

- **Connectivity matrix**:

In our work, we study the patterning on both 1-D lattice and 2-D square lattice. In later analysis, we only demonstrate two analytical solutions from two specific boundary conditions. The simulation also repeats with other types of boundary conditions, which have no simple analytical solution for eigenvalues. The MATLAB code numerically solves the eigenvalues of the connectivity matrix from given boundary conditions in simulation.

- **Dispersion relation**:

The 1-D dispersion relation, the maximum of the real part of the Jacobian eigenvalues as the function of wavenumber k, is symmetric since q(k_0_) = q(N - k_0_). In the Dispersion curve, we only show the left half, from k = 0 (isolated system) to k = N/2.

**S1 Figure: Single-morphogen patterning circuit with Gierer–Meinhardt kinetics**

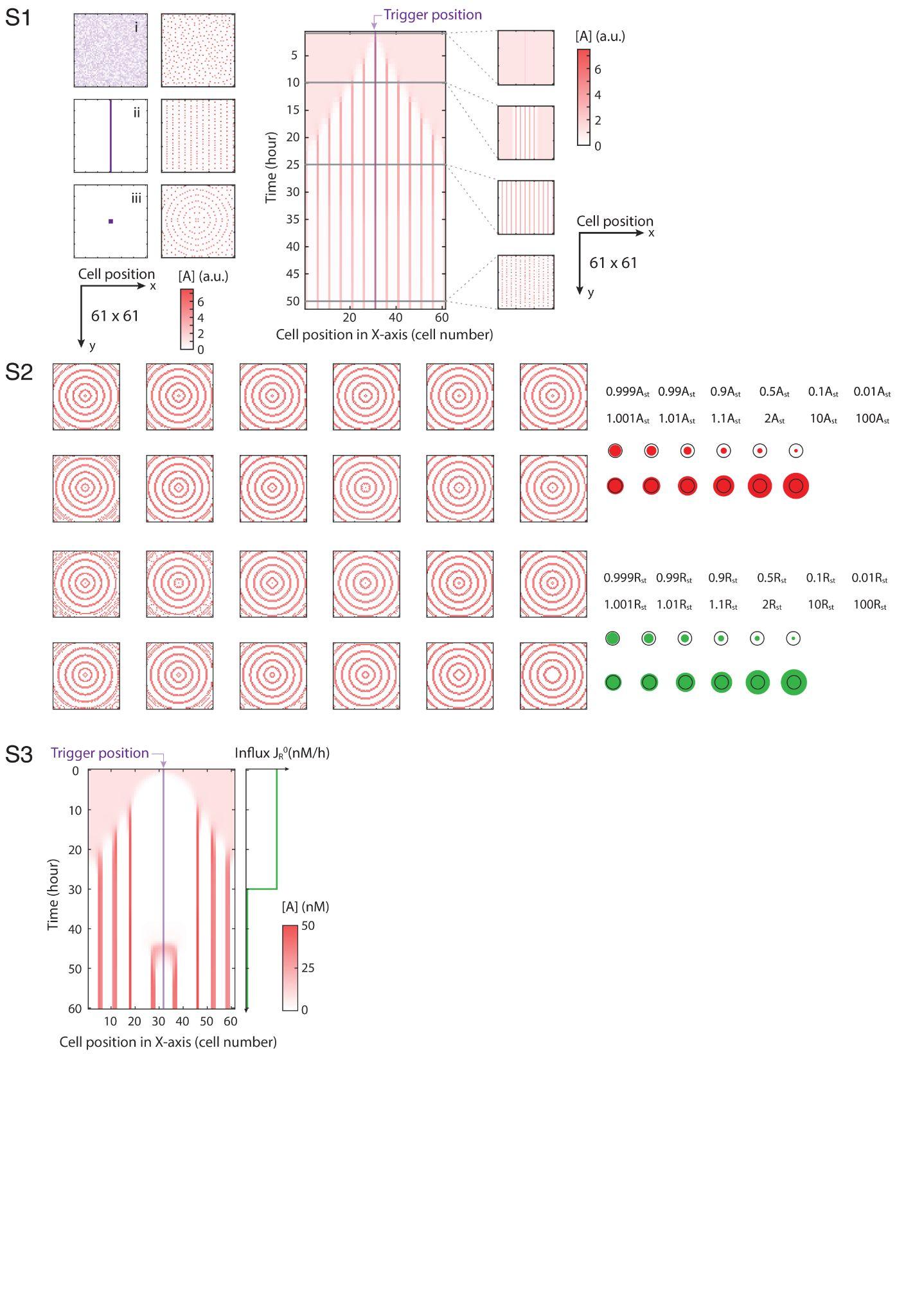

**S2 Figure: Pattern initiation is robust with a wide range of perturbations**
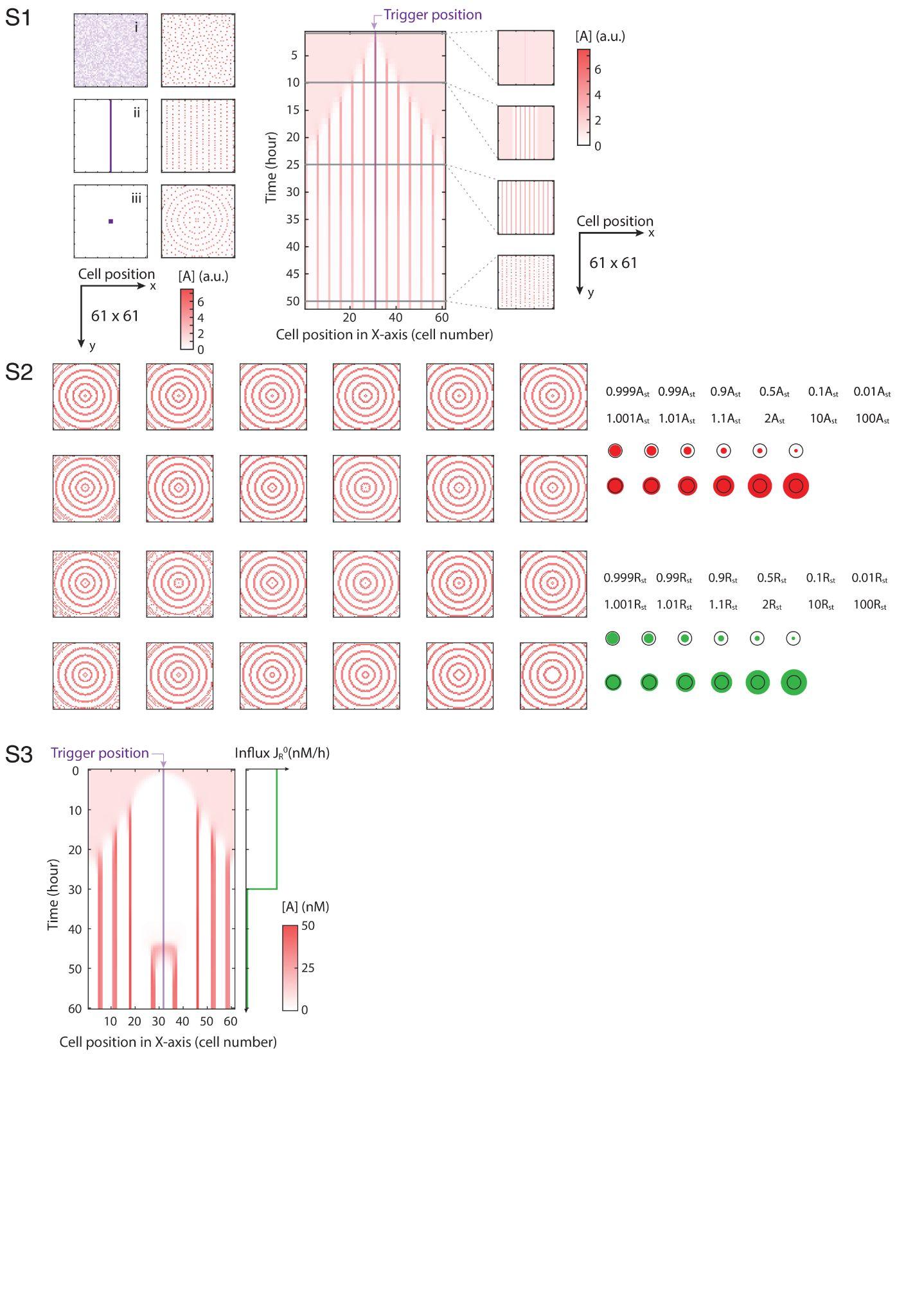

**S3 Figure: Removal of morphogen influx causes pattern stripes “fill-in” the empty space**

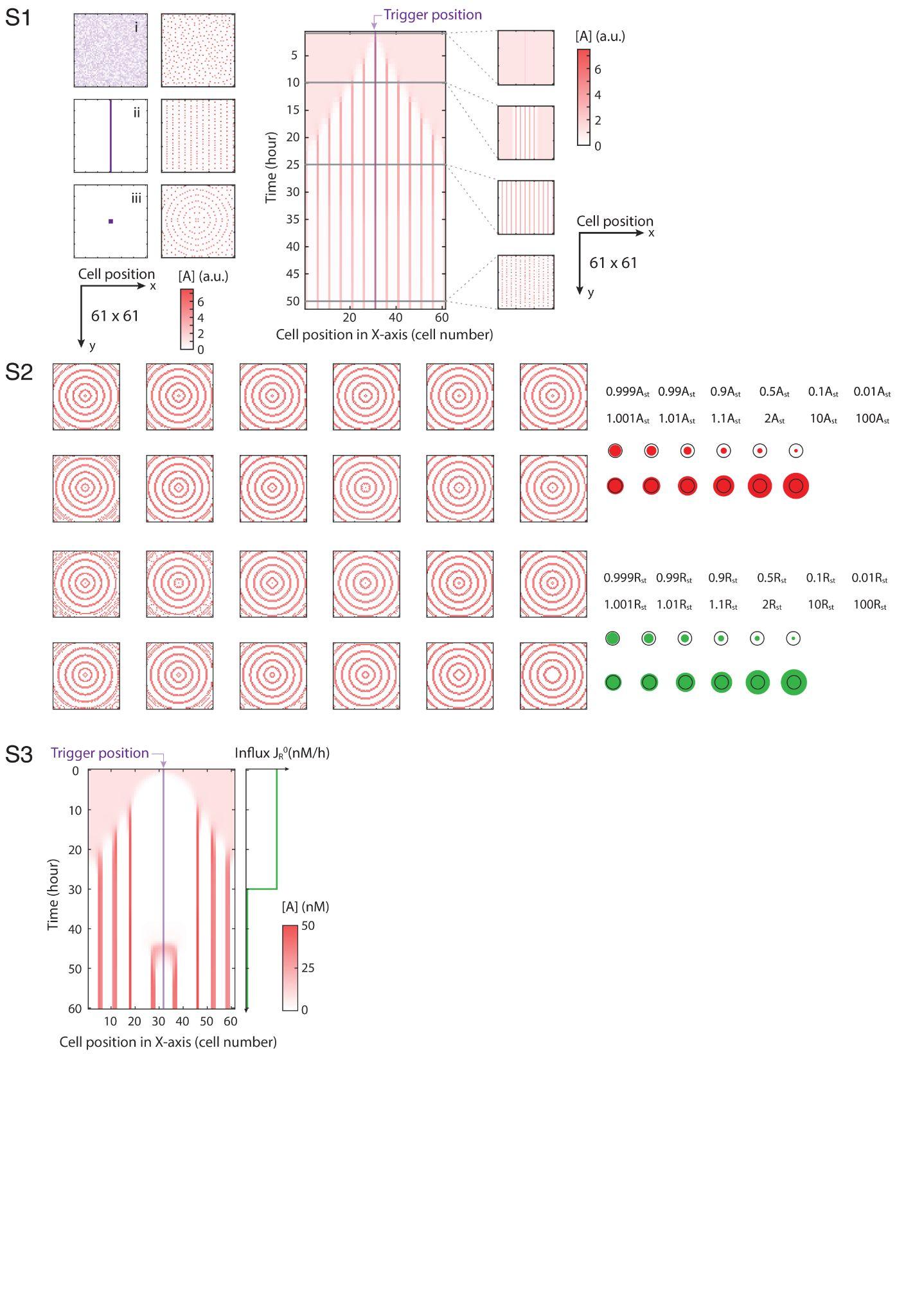

### Code reporting

Simulation code and data are accessible from CaltechDATA. Wang, S., García-Ojalvo, J., & Elowitz, M. B. (2022). Periodic spatial patterning with a single morphogen (Version 1.0) [Data set]. CaltechDATA. <https://doi.org/10.22002/D1.20060>
